# Peer-Led Leadership, Mentoring and Promotion: Conversations Among Female Academics from South Africa, Ghana and the United Kingdom

**DOI:** 10.64898/2026.07.06.736686

**Authors:** Joanna L. Elson, Marianne Venter, Phumla Sinxadi, Juliana Y. Enos, Debroah Atobrah, Gloria I. Mensah, Resia Pretorius, Sarah Guthrie, Ilse S. Pienaar

**Author notes:** These authors contributed equally: Sarah Guthrie and Ilse S. Pienaar. Correspondence concerning this article should be addressed to Dr Joanna L. Elson, The Biosciences Institute, Newcastle University, Newcastle upon Tyne, NE2 4HH, United Kingdom.

## Abstract

Here, we report a one-day, cross-cultural, peer-led workshop at the University of Stellenbosch, facilitated by academics from South Africa, Ghana and the United Kingdom. The focus was on leadership, mentoring and promotion. Using short, structured activities alongside small-group discussion, the participants were encouraged to reflect on leadership, mentoring and the perceived gap between being “ready” and being “recognised” for promotion. Descriptive survey findings and free-text reflections highlight the demand for structured peer support, reciprocal mentoring opportunities, and clearer, more transparent promotion processes. Following the event, we performed a structured review of the impact. This highlighted that the workshop participants reported that the event allowed for greater self-awareness into their own leadership approaches, a stronger commitment to purposeful mentoring, and greater confidence and renewed motivation to take concrete steps towards promotion.

## Introduction

Across universities internationally, women remain less likely to progress into senior academic and leadership roles, even when they meet promotion criteria (Murphy et al., 2021). This view was supported by a meta-analysis study showing that female academics were 17% less likely than male academics to transition to a university tenure-track position (Ceci et al., 2023). Reasons for the limitations placed on women’s career progress are rarely singular - social expectations, caring responsibilities, and uneven access to childcare or family support may restrict women’s time, flexibility and energy to engage in career-building opportunities. Despite the provision of family-friendly policies in many higher education institutions, women are still more likely to exit an academic career after parenthood than their male counterparts (Ceci et al., 2023). Within institutions, barriers may be reinforced through uneven access to mentoring and sponsorship, opaque or inconsistently applied promotion criteria, implicit bias, and workload models that may undervalue key contributions, including teaching, administration, and citizenship activities (Murphy et al., 2021). Whereas many universities have introduced initiatives to address gender inequity such as the Athena Swan Charter in the UK (https://www.advance-he.ac.uk/equality-charters/athena-swan-charter), disparities often persist between numbers of female academics in more junior roles and those attaining professorships and other academic senior leadership positions. Gender inequality may be particularly pervasive in ‘STEM’ (Science, Technology, Engineering and Mathematics), especially in traditionally male disciplines such as Physics and Engineering. The trend tends to be discipline-specific, with hiring rates into tenure-track positions often being closer to parity in fields such as Biology than in math-intensive fields such as Economics. However, for the latter disciplines, female underrepresentation may rather arise from pipeline and participation differences that occur earlier in the career path, than consistent discrimination at the hiring stage itself, though hiring outcomes for those who do apply are often relatively comparable (Williams & Ceci, 2015).

Similar barriers to career progression apply in professional and technical services career paths for women, where advancement possibilities may be even more limited due to inflexible structures and lack of formal promotion pathways (Joyce et al., 2024).

The current report describes a practical, portable intervention to improve career support for women: a one-day cross-cultural workshop designed to create a psychologically safe space for women academics and professional services/administrative staff to share experiences, compare institutional contexts, and co-develop strategies for progression. The workshop was facilitated by academics at different career stages and with wide experience of higher education contexts. The choice of peer-led facilitation supports authenticity and cultural resonance, lowers barriers to delivery, and makes the format replicable in varied institutional settings. This type of support has been highlighted as being of importance by Murphy and colleagues who showed that peer-led mentoring interventions improve academic skills, career satisfaction, and productivity (e.g. manuscripts and grants) for female faculty (Murphy et al., 2021). The structure we chose combined brief presentations and shared career stories, together with guided prompts for small-group discussions, enabling participants to move between personal reflection (i.e. what is happening in my context?) and collective sense-making (what patterns do we see, and what practical actions are available?).

## The Workshop Themes and -Design

The day-long programme integrated three inter-connected themes, namely leadership, mentoring, and promotion at work. Firstly, leadership was treated as situational, emphasising that effective approaches may vary across contexts, power dynamics and career stages. Secondly, mentoring was considered in its multiple forms, including hierarchical and reciprocal models, with attention paid to how culture and organisational structures shape access and value. Thirdly, promotion was examined as a lived process in which confidence, visibility, sponsorship, and institutional policy interact, often producing a perceived gap between ‘readiness’ and ‘recognition.’ Together, these themes were considered in a peer-led, but semi-structured context as a low-cost mechanism for both individual action planning, and advocacy to influence and reshape institutional policies and structures

In designing our workshop, we drew inspiration from our own positive experiences of professional development opportunities, and research that highlights the importance of situational and transformational leadership, where the focus is on inspiring and motivating team members towards more innovative outcomes. For example, according to a Harvard Business School study (Coronado-Maldonado & Benítez-Márquez, 2023), leaders who engage in regular self-reflection report higher emotional intelligence and decision-making capabilities. Such reflection is aided by mentoring that is appropriate to career stage and actively supported within the University. We based our approach in the belief that mentoring is valuable at all career stages and becomes especially important when women aspire to leadership positions. Nevertheless, many may struggle to cast themselves in the role of a senior leader, perceiving a conflict between empathetic treatment of colleagues, and the need to make tough decisions and require high performance from their line reports or teams. Concerns around gaining promotion, when to apply for it and how to fulfil the criteria are key concerns for academic staff, especially women. Women often wait longer between applying for promotions than their male counterparts, for example, even though they are equally successful when they do (Murphy et al., 2021). We aimed to encourage women to take stock of their skill set and to be proactive in applying for promotion when they felt ready, rather than relying on prompts from others.

## Event Overview

The event was held in-person at Stellenbosch University on the 9^th^ of September 2025. A group of colleagues from the University of Ghana joined the event online. It comprised three 90-minute workshops: *Developing Leadership Potential*, *Exploring Mentoring for Women*, and *Promotion: The Space Between Ready and Getting Recognised*. Facilitators included academics from the University of Stellenbosch, University of Cape Town and North-West University in South Africa, the University of Ghana in Ghana, and several UK universities, namely the University of Sussex, University of Birmingham and Newcastle University. There were 36 participants overall (27 from South Africa, 3 from the UK, and 3 from Ghana).

The event included presentations, insights from the individual career journeys of the facilitators, and breakout sessions with a chance to network and feedback from the small group discussions. The attendees represented a range of scientific research, technical and teaching specialisms, as well as different career stages including 20 researchers, 3 academics, 1 professor, 2 lecturers, 3 postgraduate students, 2 postdoctoral fellows, 1 senior lecturer, 1 lab manager, 1 technology transfer professional and 2 medical practitioners. The overarching goals of the workshop were to: (1) encourage reflective leadership development and adaptability across cultural contexts; (2) give insight into the value of structured, reciprocal, and cross-cultural mentoring programmes; (3) build confidence to apply for promotion and (4) recognise the diverse contributions of women to higher education, while supporting individuals’ work–life balance and assisting the navigation of personal and institutional barriers. Formalised participant feedback was gathered in an anonymised online survey sent to all registered participants, around4-weeks after the event.

The survey took the form of a link emailed to all the event participants which gave access to a Google Form, a cloud-based word processing tool forming part of the Google Workspace suite. The form could be accessed from any device with an internet connection. Anonymous data collection was ensured by altering the default settings to allow for entering data, feedback, or responses directly into the document without attaching identifying information. The invitation email to participants also clearly stated that no personal information will be shared, and this statement was repeated at the top of the Google Form. Supplementary Appendix 1 shows the sections of the Google Form presented to participants for feedback.

## Workshop 1: Developing Leadership Potential

The workshop helped participants to explore different leadership styles, understand personal preferences, and adapt approaches that would lead to greater effect. Participants completed a free https://www.teamazing.com/leadership-style-quiz/ prior to the workshop, used as a prompt for reflection, not a validated psychometric instrument. This was followed by small-group discussions based on quiz results, polls on admired leaders, and a situational leadership case study. Individuals identified their dominant leadership style by combining situational questions with behavioural preferences. The quiz distinguished between 6

Specifically, these were (1) Autocratic Leadership, involving that the leader makes decisions alone with minimal input from others; (2) Democratic Leadership that encourages team members to share ideas and take part in decision-making, and (3) Laissez-faire Leadership that gives employees a lot of freedom to make their own decisions with minimal supervision, as were described in behavioural studies by Kurt Lewin and colleagues (Lewin et al., 1939) to describe different ways in which leaders may interact with their teams. In addition, the quiz was designed to identify individuals whose management style relates to a (4) Paternalistic Leadership style, characterised by a concern for employees’ personal and professional welfare while still maintaining strong authority (Xie & Wang, 2025). Finally, the quiz allowed for identifying individuals showing tendencies towards (5) transactional leadership, where a leader motivates employees through rewards and punishments based on performance, and/or (6) Transformational Leadership that focuses on inspiring and motivating employees to achieve a shared vision and improve performance, as was identified by James Burns (Burns, 1978) and later expanded by Bernard Bass (Bass, 1985).

Participants were asked to complete a survey on leadership styles prior to the workshop. The results included the total number of participants selecting each of the given leadership styles, as either primary or secondary (Figure 1): Paternalistic (13 primary, 6 secondary) and Democratic (10 primary, 11 secondary) styles were frequently selected, with fewer individuals selecting Transformational (2 primary, 3 secondary), Autocratic (0 primary, 3 secondary) and Transactional (0 primary, 3 secondary) styles; none conformed to a Laissez-faire leadership style (0 primary, 0 secondary).

**Figure 1.**
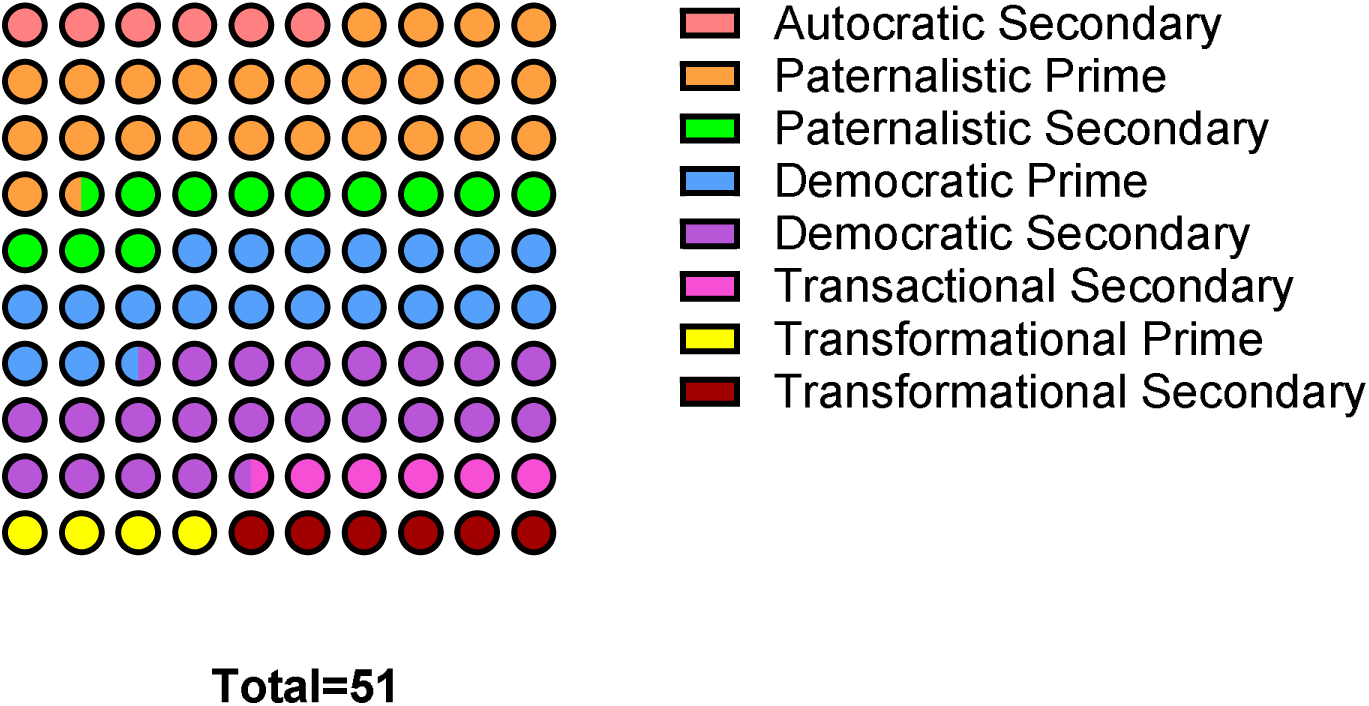
Distribution of self-reported primary (solid bars) and secondary (hatched bars) leadership styles selected by workshop participants from a pre-event online quiz. Paternalistic (13 primary, 6 secondary) and Democratic Leadership (10 primary, 11 secondary) styles were most frequently endorsed, reflecting preferences for supportive direction combined with inclusive decision-making. Transformational (2 primary, 3 secondary), Autocratic (0 primary, 3 secondary), and Transactional Leadership (0 primary, 3 secondary) styles were less common, with no participants conforming to the Laissez-faire Leadership style category (0 primary, 0 secondary). Data shown were derived from anonymous post-event survey responses (n=36 participants total).

These responses indicate that participants favoured a mix of paternalistic and democratic styles. Whereas the term *paternalistic* can feel like a misnomer in the context of women’s leadership, participants appeared to use it to signal a *protective, responsible* stance: Making decisions in the interests of the group and providing clear direction when needed. However, they also noted that if over-used, this approach can limit autonomy and slow the development of others, creating dependency rather than capability. A *democratic* style was widely favoured, with participants emphasising inclusive decision-making: actively seeking input, building consensus where possible, and ensuring that team members’ perspectives were heard and incorporated into strategy and planning

Participants also acknowledged the practical trade-off that democratic approaches can be slower and are not always feasible in time-critical situations, requiring occasional shifts toward more directive decision-making. Democratic approaches may reflect inclusive leadership values and a culture of listening to others; however, participants also noted that prolonged consensus-seeking can sometimes act as a form of risk avoidance, especially when confidence or authority is limited.

We note that attendees represented a wide range of roles and career stages. raising the possibility that the apparent preference for paternalistic and democratic styles may partly reflect role-related expectations and limits on decision latitude that are context-specific. In the follow-up survey free-text responses, most participants (6/7) described active or planned changes in their own leadership practice, often moving from paternalistic oversight towards more democratic, structured delegation and wider consultation, consistent with a shift toward participative/inclusive leadership.

There was discussion of how different types of leadership style might be required in different contexts. There was also a discussion on the balancing of support versus dependency, allowing space for team members to develop their own independence. The need to provide clear direction for the team could sometimes be problematic for women operating in very male-dominated workplaces. A transformational style was seen as important in a visionary context or for change management. Two respondents explicitly framed an intentional move toward transformational practice in specific contexts (e.g. student-led facilitation), alongside heightened self-awareness and a sustained commitment to reflective adaptation of style over time.

Quotes from participants showed the value of reflection within the session, while highlighting the issues that emerge in choosing leadership styles:

Quote 1**: ‘***Developing Leadership Potential increased my self-awareness of the style(s) I use and how to improve and balance other styles.’*
Quote 2: *‘Yes, I am trying to be a bit less paternalistic and encourage my team members to take on more leadership so that they can develop their skills independently e.g. through delegation of tasks, without me being overly concerned about their wellbeing all the time.’*
Quote 3: ***‘****Considered changing to be more transformational leadership style, especially in my facilitation sessions with student groups - as the format is naturally set up for students to be leading the session.’*

Overall, participants acknowledged the need to adapt leadership to the situation, reflecting on and learning from the success of different approaches, having to accept less than optimal outcomes on occasion, and to move forward from these.

## Workshop 2: Exploring Mentoring for Women

This session presented insights from the University of Sussex – University of Ghana project entitled *A Cross-cultural Reciprocal Mentoring Scheme*, funded by the British Council, and led by Professors Sarah Guthrie (Sussex), Deborah Atobrah (Ghana) and Dr Juliana Enos (Ghana). This scheme provides an alternative to traditional, hierarchical mentoring, as a peer-to-peer mentoring opportunity, and recognising the role of cultural context in women’s lives and career progression. In the session, Prof Sarah Guthrie and Dr Juliana Enos (Ghana) explained the scheme - a year-long programme in which around 23 women from each University (46 total) formed pairs to undertake a structured mentoring journey. Training sessions, both jointly online and in person at each University formed the background, and importantly, developed cultural competency and listening skills in the participants, The women were encouraged to think about their expected benefits from the scheme, and to actively commit to the partnership.

Mentoring pairs then met six times over six months with a session structure of topics with prompt questions for each meeting. Online drop-ins were provided each month at which the project team talked to participants about their experience and offered support where needed. The project gathered feedback using focus groups and questionnaires, and the project teams undertook reciprocal visits to the UK and Ghana, to collaborate on various outputs, including a career support toolkit.

Two key aspects of the mentoring scheme have been its reciprocal and cross-cultural nature. In the context of the Stellenbosch workshop, a framework was presented for understanding different types of mentoring, for example hierarchical, reverse mentoring or reciprocal mentoring. Whereas traditional hierarchical mentor (senior colleague as mentor, junior colleague as mentee) can be useful, it sometimes reinforces existing inflexible structures within the workplace and embeds conformity. Reciprocal mentoring, as implemented here, emphasises peer learning. The added cross-cultural context was also of relevance to the workshop at Stellenbosch, which included white women and women of colour. Issues of diversity are key to understand the intersectionality and added barriers that impede women’s career advancement in higher education in Africa and the UK. The cross-cultural mentoring programme allowed women to share lived experience and understand each other’s contexts, challenges, and approaches for success. Some generalities about mentoring could be drawn from the experience of running this mentoring scheme, including the need for active support and help to maintain mentoring relationships over time. Not all mentoring relationships were successful and sometimes when partnerships could not continue, alternative mentoring partners were found. We also noted technological and workload challenges in conducting a collaboration across continents. However, there was overall very high enthusiasm for the scheme; many had formed lasting friendships and/or developed research or education collaborations or other types of partnership. Mentoring meetings were also seen as a psychologically safe space where personal and professional issues of importance to the individual could be discussed in a non-judgemental way.

During the Stellenbosch workshop, participants interrogated their own experiences of mentoring, whether they had been given such opportunities, and how useful this had been for their development and career progression. They were also asked to consider the different types of mentoring, hierarchical, reciprocal or reverse mentoring, and how these formats might work in specific contexts. Most participants fed back that their own mentoring opportunities to date had been very limited, until they reached senior levels. Some examples of support were given, e.g. Stellenbosch University’s SUNRISE (Stellenbosch University Research & Innovation Strategic Excellence; https://www.su.ac.za/en/node/865 programme, a selective two-year development programme aimed at high-performing mid-career academics, providing mentorship, personalised academic support, coaching, and protected time for research. Other South African initiatives for mentorship support given to academics that were cited by the participants included South Africa’s Department of Higher Education and Training’ ‘Future Professors’ Programme (https://futureprofessorsprogramme.co.za/). Informal peer to peer mentoring and life coaches were seen to provide crucial support where available. However, activities such as mentoring, and professional development in general, were often seen by the participating cohort to be of a low priority when workload is heavy, and many demands are placed on time.

Other perceived barriers included a lack of mentors, lack of a formal process to ensure follow-through where mentorship programmes were in place, overbearing line managers who left little time for personal or professional development and a lack of transparency about job expectations. Other barriers to undertaking mentoring were seen to be culturally specific. Ghanaian colleagues mentioned that extended families sometimes contribute to childcare but on the other hand produce expectations of women being caregivers for wider networks of relatives. In other cultures, the nuclear family puts the onus on the life-partner to provide support or means seeking organised childcare. Overall, there was a strong desire for more mentoring opportunities, including ones that were more tailored to particular career stages, and especially aimed at early career researchers. In the post-event reflections, most respondents described concrete changes or intentions to change their mentoring practice, suggesting acknowledging the value of mentoring, and making time for its practice.

Among respondents, 4/5 (80%) reported that they had either already begun, or intended, to expand or adapt their mentoring relationships, whereas 1/5 (20%) indicated that time pressures had so far prevented any change. Several responses showed explicit intent to mentor early-career colleagues, including across disciplinary boundaries. Notably, one participant described asking their PhD supervisor to take on a clearer mentoring role through monthly meetings, suggesting that even relatively basic expectations of developmental support are not consistently in place. This may reflect a need for women to ‘manage up’ by making their mentoring and career support needs explicit to their managers, rather than waiting for this support to be forthcoming. Another respondent called for stronger mentoring of postdoctoral researchers and junior lecturers, particularly to improve the quality, rather than simply the quantity, of papers and student outputs. Taken together, these responses indicate a clear appetite for more structured, purposeful mentoring, especially at earlier career stages.

Reported changes in mentoring practice included more regular and intentional check-ins, mentoring tailored to individual needs, and greater emphasis on balancing professional development with wellbeing. Participants also described greater empathy and patience, clearer expectations around roles and responsibilities, and more proactive strategies, such as gentle “nudges”, to encourage mentee engagement. Several comments further reinforced the importance of informal peer-mentoring cultures within departments, particularly where formal mentoring schemes are unavailable or otherwise limited.

Quote 1: “*Since the workshop, I’ve been more intentional about tailoring my mentoring to individual needs. For example, I now check in more regularly with students and early-career colleagues, actively seek their perspectives on challenges, and provide guidance that balances career development with personal well-being.*”
Quote 2: “*I am not in a formal mentoring role, but mentor students and colleagues in an informal way. I do not really think I can make these interactions more efficient, except for maybe trying to encourage more informal interactions in the department.”*
Quote 3: *“I wish more was done to mentor postdocs and junior lecturers at* [*] *for quality of papers and not just quantity - happy to get involved.”* * University name redacted to ensure participant anonymity.

Responses indicated a clear direction of travel toward structured, responsive mentoring that builds confidence through trust, clarity, and consistency, while recognising practical constraints (particularly time) and the importance of cultural and organisational context.

## Workshop 3: Promotion - The Space Between Ready and Getting Recognised

This workshop examined why women are less likely to apply for promotion even when they may have an appropriate track record and have fulfilled relevant criteria. A recent article in Nature Communications showed that there is a gender difference between the level of self-promotion using social media channels, with women around 28% less likely than men to self-promote (Peng et al., 2025). Participants heard personal stories, including the different career journeys of three academics from South Africa and UK who reflected on barriers, and identified practical actions. These stories presented instances where “invisible work” that contributed to the progress of a department was not recognised in the promotional process that only considers measurable outcomes such as number of papers; where reasons for an unsuccessful promotion application seemed arbitrary such as “not enough papers” or “not famous enough” while these criteria could not be clearly defined upon request; and where a successful promotion was seen in hindsight as long overdue, indicating a lack of institutional interest in the promotional journeys of women.

Individual career reflections included a common theme that women are often expected to fulfil all criteria of promotion before applying and wait a long time before applying. Sometimes individuals are reticent to put themselves forward without their line-manager’s support, even though they may be successful if they do. The systemic issue that academic promotion is usually awarded once one is already doing the job that is being applied for, was alluded to. This framing strongly aligned with participants’ subsequent survey comments, which emphasised both “internal” barriers (confidence, self-advocacy) and “external” barriers (visibility, access to opportunities, workload and institutional culture).

Participants were asked to make loose word associations with the term ‘Promotion’ and mentioned the following: *Bureaucracy, Recognition, Intimidation, Growth, Work, Status, Power, Prestige, Paperwork, Accountability, Committee meetings, Imposter syndrome, Responsibility, Commitment, Reflection.* Ideas of imposter syndrome chimed with comments that were made around being told a person was ‘not famous enough’ to be promoted. Despite the impediments it was seen as crucial to increase the number of female role models among senior faculty in universities, to inspire others. Notably, the associations captured both aspiration (growth, status, recognition) and friction (bureaucracy, intimidation, paperwork), echoing the “space between ready and recognised” theme.

Practical and systemic barriers to promotion in the various institutions were perceived to be non-standard criteria, lack of transparency in who got promoted, penalisation of part-time working and caring responsibilities, financial limitations of the university, lack of support from line managers and favouritism. Suggestions for how to plan your promotion applications were to keep detailed records, request regular 1:1s with line managers, manage workload strategically, set career development plans and leverage networks. In the survey responses (5 in total), barriers clustered across structure/policy (e.g. NRF expectations and narrow definitions of “focus”), workload/time pressure, recognition of non-linear trajectories (including part-time working), departmental culture/leadership, and confidence/self-advocacy. Several respondents listed multiple barriers, and no single issue dominated.

Participants also identified that high-leverage institutional supports would make the biggest difference: funded postgraduate places to build research capacity; clearer and more transparent promotion pathways; structured mentoring and senior sponsorship; meaningful administrative relief; and research forums for brainstorming and strategy.

Alongside leadership/professional development opportunities, policies that protect work–life balance (flexible scheduling and equitable allocation of administrative responsibilities) were seen as critical to reaching career potential.

Quote 1: “*You essentially need NRF* [The National Research Foundation, a government agency that supports and promotes research and innovation across all fields of knowledge in the country] *rating and the NRF wants to see a clear focus in your research… Applying for promotion has a hard time to get to the top of the priority list.”*
Quote 2: “*Clearer career pathways, less admin.*”

## Discussion

Women remain under-represented in senior academic roles, not simply because of individual confidence or ambition, but because of a complex interplay of personal and institutional factors. For women to identify these factors and think through what mitigations and proactive approaches to support their career advancement is possible, space and time is needed for personal reflection. Our workshop was structured to provide this non-judgemental space with peers, where hearing from others can bring into focus an individual’s circumstances, and prompt planning for the future (Chan, 2013). The findings from our leadership workshop demonstrated women’s preference for Paternalistic and/or Democratic Leadership styles. This pattern is broadly consistent with the wider leadership literature: Women leaders, on average, tend to be somewhat more participative/democratic and also more transformational in style, while men are reported to be more likely to show laissez-faire and some transactional “management-by-exception” behaviours (Eagly et al., 2003). We did not observe clear cross-cultural differences in preferred leadership styles within this group. That is compatible with cross-national evidence suggesting that gender differences in leadership behaviour are often modest and context-dependent and may vary by societal setting rather than showing a simple, uniform pattern across countries (van Emmerik et al., 2008). Nonetheless, given the shared professional context and modest follow-up numbers, we can say that no systematic differences were observed in this sample, rather than asserting that cross-cultural differences do not exist. Snaebjornsson and colleagues argue that leaders often model their conduct in line with genderless societal expectations of leadership, while followers may still hold gendered expectations, with associated social differences creating scope for perceptual dissonance when women adopt more “masculine-coded” styles (Snaebjornsson et al., 2015).

Mentoring discussions highlighted a parallel theme: where formal mentoring is limited, informal peer support becomes essential, yet time pressure, unclear expectations, and overbearing supervision can restrict access to developmental opportunities (Mousa et al., 2023) with other studies noting the complexity of the issues and questioning the overarching value of traditional models (House et al., 2021).

Our discussions highlighted the idea that hierarchical mentoring may reinforce existing institutional structures, and patterns of access to opportunities sponsorship, and advancement. It may become self-perpetuating of existing inequalities, particularly where mentoring is unevenly available, dependent on senior goodwill, or embedded within opaque academic cultures (Murphy et al., 2021). By contrast, we suggest that reciprocal mentoring models offer an alternative that values peer learning, recognises cultural context and difference, and creates space for more candid discussion of shared barriers and practical strategies for progression (Levy-Tzedek et al., 2018; Pfund et al., 2022). International and cross-cultural mentoring can also enhance international opportunities and networking for participants, bolster University partnerships and improve understanding of how to work and lead within diverse teams and communities of colleagues. In this case, prior connections between some online participants and facilitators appeared to support discussion and continuity, suggesting that cross-cultural initiatives may work best when relationship-building begins before the event and is reinforced through planned follow-up (House et al., 2021; Mousa et al., 2023). The Workshop 2 feedback highlighted when formal mentoring is limited, women often rely on informal peer support, yet access to even this support is shaped by time pressure, local culture, and organisational hierarchy.

Comments suggested that developmental support is often sustained through informal departmental relationships whereas it is preferable that institutions should create conditions in which mentoring, visibility, and development are actively enabled rather than left to chance (Mousa et al., 2023). As reflected in the systematic review by House and colleagues, the evidence base for the effectiveness of mentoring is heterogeneous, and traditional hierarchical models do not consistently overcome the wider structural barriers women face (House et al., 2021). Discussions concerning workshop 3 tied these strands together. Participants’ word associations captured both aspiration and friction with reported barriers clustered around workload, opaque criteria, limited visibility and sponsorship, and uneven recognition of non-linear trajectories (including part-time work and caring). The generally held opinion that many women apply for promotion too late may map onto recent evidence that gender gaps in progression persist even when accounting for research outputs, pointing to unequal returns and institutional constraints rather than individual ambition as the sole explanation (Czech et al., 2024).

The call for “clearer career pathways, less admin” therefore reflects a practical theory of change: Make criteria transparent and apply them consistently, protect time by reducing administrative/service overload, and pair mentoring with genuine sponsorship that increases access to high-visibility opportunities (Schwartz et al., 2024). Overall, the event demonstrates a portable model: Universities can create effective peer-led development spaces without specialist facilitation, using a simple structure, protected time, and cross-role participation to generate both individual actions and institutionally relevant insights. Although such small, focused workshops are not likely to result in changes at an institutional level, feedback from the participants highlighted the impact of such events at the individual level. Participants indicated the value they found from attending the event in the anonymised post-event survey.

The responses indicated an increased awareness of their own leadership styles and those of others, and how small changes in leadership style may improve the chances of attaining desired group outcomes. Additionally, participants noted the impact on their own perceptions of hearing about the experiences of peers. Lastly, while participants acknowledged and recognized the universal barriers to career development, a consensus was reached on the importance of implementing change on an individual level, especially for those in positions to mentor and support women in more junior positions (i.e. ‘sending the elevator back down’). The workshop therefore afforded participants the opportunity to reflect on these issues and their roles in the wider context of university culture.

## Conclusion

This peer-led, cross-cultural workshop demonstrates that structured, low-cost interventions can effectively foster self-awareness in leadership styles, promote reciprocal mentoring practices, and bridge the confidence gap to promotion for women academics, yielding immediate individual impacts like intentional behavioural changes despite persistent institutional barriers. Ultimately, such initiatives empower women to enact personal agency while advocating for systemic reforms like transparent criteria and reduced administrative burdens. Future efforts should prioritize peer facilitation across diverse cultural contexts, integrate pre- and post-event surveys for measurable outcomes, and scale through hybrid formats with sustained follow-up networks to amplify long-term career progression.

## Ethics statement

Ethical clearance for including participant feedback and observational notes on the group exercises were obtained through the University of Birmingham’s ethics committee (reference number: ERN_5502-Dec2025).

## Acknowledgements

The event was part-funded by a University of Birmingham ‘International Engagement Fund’ grant, awarded to Dr Ilse S. Pienaar. The Cross-cultural Reciprocal Mentoring scheme is funded by a British Council ‘Going Global Gender Equality Partnerships’ grant, awarded to Professors Sarah Guthrie and Debroah Atobrah, and Drs Juliana Enos and Gloria Mensah.

## Appendix 1

## Workshop 1: Developing Leadership Potential (leadership styles)

### Question: Since the workshop, have you changed, or are you considering changing your academic leadership style?

- “Considered changing to be more transformational leadership style, especially in my facilitation sessions with student groups - as the format is naturally set up for students to be leading the session.”
- “I am slowly learning to be less paternalistic and do things for people, to being democratic and let the people do the things.”
- “Yes, I am trying to be a bit less paternalistic and encourage my team members to take on more leadership within the team so that they can develop their skills independently e.g. through delegation of tasks, without me being overly concerned about their wellbeing all the time.”
- “Increases self-awareness of my leadership styles”
- “Yes, to diversity my leadership style more”
- “Planning to be more transformational. It requires some thinking about how to position myself self and what to do differently.”
- “Since the workshop, I’ve been more intentional about collaborative and inclusive leadership. For example, I now actively seek input from all team members in decision-making and balance task-focused goals with attention to mentorship and team well-being. I plan to continue reflecting and adapting my style as needed.”

## Workshop 2: Exploring Mentoring for Women (mentoring practice + mentoring relationships)

### Question: Since the workshop, have you changed, or are you considering changing your mentoring practices?

- “Yes, really thought deeply about making a concerted effort in my role as a mentee and being more open about my personal journey to benefit more from the mentor. Also, I’m struggling with being a mentor (to a high school learner) but from after the workshop I tried to subtly nudge the mentee to be more engaged - on the plus side I feel less anxious and frustrated but I’m not sure the mentee has gained much from our relationship.”
- “I am not in a formal mentoring role, but mentor students and colleagues in an informal way. I do not really think I can make these interactions more efficient, except for maybe trying to encourage more informal interactions in the department.”
- “To be more patient and also put myself in the mentee’s shoes”
- “Not really, no”
- “Aim for well-established responsibilities for each team member”
- “Yes, I have changed it to something that suits me better.”
- “Yes. To be more engaging and focus on influencing my mentees lives as best as I can.”
- “Since the workshop, I’ve been more intentional about tailoring my mentoring to individual needs. For example, I now check in more regularly with students and early-career colleagues, actively seek their perspectives on challenges, and provide guidance that balances career development with personal well-being.”

### Question: Since the workshop, have you, or do you intend to pursue new mentoring relationships or adapt existing ones?

- “I did not really have time to think about this. I have been super busy.”
- “Yes”
- “I have indirectly asked my PhD supervisor to take on a bit more of mentor role for me by meeting with me monthly to explore and expand on new research ideas.”
- “I wish more was done to mentor postdocs and junior lecturers at ** for quality of papers & students and not just quantity - happy to get involved”
- “Since the workshop, I have been more intentional about expanding and adapting my mentoring relationships. I am actively seeking opportunities to mentor early-career colleagues across disciplines and ensuring that existing mentees receive guidance tailored to their individual goals and challenges.”

### Question: Have cross-cultural or peer connections formed at the workshop persisted or expanded?

- “Yes, it was good to chat to students in Physiology. I had contact afterwards. Maybe we can do something together on radio-therapeutic isotopes in micro-clots.”
- “No”
- “No, I haven’t connected with anyone again after the workshop”
- “Met lots of new people which was great but have not had follow up connections (yet!).”
- “Not really, and I’m a bit disappointed about that. I would really value staying in touch with the women I met at the workshop and hope there might be opportunities to connect or collaborate in the future.”

## Workshop 3: Promotion: The Space Between Ready and Getting Recognised Question: What barriers to promotion remain most noticeable for you?

- “You essentially need NRF rating and the NRF wants to see a clear focus in your research. My research is over different fields with applications from astrophysics, medical isotopes, quantum technology to scholarship of teaching and learning. I know one can apply for promotion without the formal NRF rating, but you will probably still be evaluated on the same scale. I have worked part time for many years, so I should maybe test the system and find out whether that will be taken into account in evaluation for promotion. But my main problem is that I am too busy with activities that are important and make a difference. Applying for promotion has a hard time to get to the top of the priority list.”
- “Self-confidence and overall emotional intelligence”
- “Putting myself forward and feeling like I have the credentials (I have not had a straightforward academic trajectory) to be promoted”
- “Weak leadership in departments”
- “The most noticeable barriers to promotion for me remain the limited visibility of women in senior academic roles, unequal access to high-profile leadership opportunities, and the persistent challenge of balancing research, teaching, and administrative responsibilities with personal and family commitments.”

### Question: What institutional supports do you feel would make the biggest difference to help you attain your career potential?

- “Bursaries for deserving postgraduate students.”
- “Clearer career pathways, less admin”
- “More support from senior colleagues who have had similar experience”
- “More dynamic scientific circles to brainstorm and strategize. also more admin support (in both teaching and research).”
- “Institutional supports that would make the biggest difference include structured mentorship programs, transparent criteria and pathways for promotion, access to leadership and professional development opportunities, and policies that support work-life balance, such as flexible scheduling and equitable distribution of administrative responsibilities.”

## Event-wide evaluation (across workshops)

### Question: Do you feel more confident or equipped to pursue career advancement opportunities as a result of having attended the workshop?

- “4 – Yes”

### Question: What was your most valuable personal takeaway from the workshop event?

- “It made me think that I should maybe consider applying for promotion. At least the thought made it onto the bottom of my priority list.”
- “The experience overall was valuable to me - hearing the stories of the other ladies including the facilitators”
- “Learning more about my leadership style and hearing others’ experiences around promotion”
- “For me 1. Developing Leadership Potential increased my self-awareness of the style(s) I use and how to improve and balance other styles”
- “My most valuable takeaway was the importance of intentional, inclusive leadership and mentorship, and how small, conscious changes in approach can have a meaningful impact on team dynamics and career development. This relates most directly to Workshop 1: Developing Leadership Potential.” Inclusive leadership and mentorship -

## Notes

### Competing Interest Statement

The authors have declared no competing interest.

